# Interference competition influences virulence in *Metarhizium* coinfections

**DOI:** 10.64898/2026.10.02.756158

**Authors:** Najmussher Ghani, Cemil Ögüt, Naomi Kimenyi, Sophie Kilker, Charlotte Rafaluk

**Affiliations:** Evolutionary Biology, Institute of Biology, Freie Universität Berlin, Königin-Luise-Straße 1-3, 14195, Berlin, Germany; Fakultät für Biologie und Vorklinische Medizin, Universität Regensburg, Universitätsstraße 31 D-93053, Regensburg

**Author notes:** Corresponding author name and email: Charlotte Rafaluk. Equal contribution (Najmussher Ghani, Cemil Ögüt).

**Keywords:** Coinfection, insect, fungus, host-parasite interaction, virulence evolution, coevolution, trade-off hypothesis

## Abstract

Coinfections are common in a nature and often involve competition among coinfecting parasites to extract resources from the host. As a consequence of this competition, parasites can face trade-offs between investing in competition with other parasites and exploiting the host for their own growth and transmission. Such within-host interactions among parasites can either increase or decrease virulence depending on how they alter parasite density, resource use, and host damage. We tested the inhibitory effect of different species of the entomopathogenic fungus *Metarhizium* on one another *in vitro* and the impact of coinfection on virulence towards *Tribolium castaneum*. We found that *Metarhizium pemphigi* is able to inhibit all other isolates of *Metarhizium* tested, whereas *Metarhizium pingshaense* was relatively resistant to inhibition by other *Metarhizium* strains. Virulence towards the host, *Tribolium castaneum*, differed very little among isolates tested. We did, however, observe differences between virulence in single and coinfections in two cases. Where there were virulence differences, these were both consistent with *in vitro* inhibition patterns and interestingly resulted in reduced virulence in coinfection. This is inconsistent with broader patterns of virulence in coinfection, where virulence seems to be driven by the more virulent parasite. It is, however, consistent with data from other systems where there is evidence of interference competition. Our data thus support the idea that, in contrast to other forms of between parasite interaction, interference competition results in reduced virulence in coinfection.

## Introduction

Coinfections are both common in nature and are ecologically more complex than single infections (Zilio & Koella, 2020). In coinfection, parasites interact directly with the host and also with one another (Betts *et al*., 2016). Whereas in many single infections parasite virulence is predicted to optimise at an intermediate level (Anderson & May, 1982; Leggett *et al*., 2013), predicting the effects of coinfection on virulence and parasite fitness is not straightforward (Bose *et al*., 2016). Research on the impact of coinfection on the overall virulence of infections is increasing (Venter *et al*., 2022), with outcomes dependent on a wide range of ecological factors (Alizon *et al*., 2013; Milutinović *et al*., 2020; Seabloom *et al*., 2015). A recent meta-analysis has shown that in general the most virulent parasite in coinfection tends to determine overall virulence, but with substantial variation in outcomes (Rafaluk *et al*., 2026).

The fitness of individual parasite genotypes in coinfection is highly dependent on the within host replication rate (Balmer *et al*., 2009; Bell *et al*., 2006; De Roode *et al*., 2005) and the pattern of host exploitation used by the other coinfecting parasite (Ben-Ami *et al*., 2011; Frank, 1996), as well as the nature of the interaction between coinfecting parasites (Alizon & Lion, 2011; Massey *et al*., 2004; Mideo, 2009). These interactions can differ from extremely positive to extremely negative and range from highly competitive e.g. (Garbutt *et al*., 2011) to co-operative e.g. (Buckling *et al*., 2007). Relatedness among co-infecting parasites is often a key factor determining the type of interaction occurring between coinfecting species (Frank, 1992; Gleichsner & Minchella, 2014), although many other ecological factors play a role (Gao *et al*., 2016; Kinnula *et al*., 2017; Tracy *et al*., 2021).

If infecting parasites secrete public goods which contribute to virulence, less virulent parasites, which do not themselves secrete as large quantities of a given public good, may benefit from the presence of a coinfecting parasite that contributes more, resulting in reduced virulence in mixed infections (Buckling *et al*., 2007). Alternatively, when host resources are limited, parasites may invest in competition (Frank, 1996; Massey *et al*., 2004). Competition may be exploitative, where parasites compete indirectly for a limited host resource (De Roode *et al*., 2005; Mideo, 2009); apparent, where one parasite evokes an immune response which negatively impacts the competitor (Clay & Rudolf, 2019; Råberg *et al*., 2006; Read & Taylor, 2001) or direct, acting via mechanisms such as interference competition where coinfecting parasites directly inhibit one another (Gardner *et al*., 2004; Massey *et al*., 2004).

Entomopathogenic fungi are particularly likely to find themselves in coinfections. Many are generalists (Hajek & St. Leger, 1994; Wang & Wang, 2017), thus likely to find themselves in a broad range of hosts, potentially bringing with it exposure to a broad range of other parasites. Additionally, entomopathogenic fungi produce long lived conidia (Hong *et al*., 1999, 2001), meaning that multiple fungal genotypes are likely to be present in the environment at the same time. We hypothesised that entomopathogenic fungi of the genus *Metarhizium* may have evolved strategies to maintain fitness in coinfection, impacting the virulence of coinfections. *Metarhizium spp*. are generalist pathogens of arthropods (Milutinović *et al*., 2020; Zhao *et al*., 2023), with applied relevance for biocontrol of *T. castaneum* (Peng *et al*., 2022; St. Leger & Wang, 2020; Velavan *et al*., 2022) and thus of particularly broad relevance. Indeed, coinfections of *Metarhizium anisopliae* and *Metarhizium robertsii* result in lower virulence compared to single species infections, as a consequence of within host competitive exclusion (Sheng & St. Leger, 2024). We tested whether isolates of *Metarhizium* with differing degrees of phylogenetic relatedness inhibited one another *in vitro* and how coinfections of different strain pairs differed in virulence from their respective single infections towards the model host *Tribolium castaneum*.

## Materials and Methods

### Host

The *T. casteneum* population Cro1 (Milutinović *et al*., 2013), was used for all experiments. Adult beetles were reared on organic wheat flour (type 405) containing 5% dried yeast at 25±1° C and 70±5% relative humidity. The temperature of 25±1° C was chosen as it has been used previously for the rearing of *T. castaneum* with no impact on survival (Al-Zurfi *et al*., 2023), and because *Metarhizium* induced mortality was too rapid and the mortality window too short, to compare at the standard rearing temperature of 30±2^0^C. Prior to the experiments described here, we conducted preliminary survival assays on different host life stages and at different temperatures to find the best conditions for our experiment. We did not observe any infection in adult beetles even at a dose of 1 x 10^9^ spores/ml at temperatures ranging from 25-33° C. When testing infection in larvae, we observed frequent cannibalism. Pupae are both susceptible to infection and do not present the complication of cannibalism in age-synchronised populations. Consequently, 0 to 1-day-old pupae were used in all our *in vivo* experiments.

### Parasites

We used a total of five isolates of *Metarhizium*, belonging to four species: *Metarhizium lepidiotae* ARSEF 8064, *Metarhizium pingshaense* ARSEF 8092, *Metarhizium pemphigi* ARSEF 9358 (Zitlalpopoca-Hernandez *et al*., 2022) and *Metarhizum brunneum* strains *KVL* 13-06 and KVL 16-36 (Steinwender *et al*., 2014). Fungal isolates were grown on 2.4% PDA (Potato-Dextrose-Agar) agar plates with 2% at 20° C. Spores were harvested after 14 days using 0.01% Tween 80 as a surfactant.

### Radial growth

A spore suspension of 1×10^7^ spores/ml was prepared in 0.01% Tween 80. For each *Metarhizium* isolate tested, 10µl of 1×10^7^ spore/ml suspension was pipetted onto a 9cm PDA plate. Inhibitory effects were tested between all isolate pairs fully factorially including self-inhibition (i.e., two drops of the same isolate suspension). To test for inhibitory effects, two droplets were placed 2 cm apart on a 9cm PDA plate and the plates incubated at 20°C. The diameters of both spots were measured after 7 days. For each run of the assay, each isolate was also grown alone on PDA plates, with three replicates, to provide a measure of baseline radial growth. The colonies all had roughly spherical growth (example in Figure S1) and thus the horizontal diameter were measured. Both colonies on each plate were measured and the growth in combination with a challenger subtracted from the growth rate alone, to create a measure of inhibition. The experiment was repeated three times, with five replicates of each isolate combination per run.

### Parasite-induced host mortality

Pairs of isolates were selected in which to test parasite-induced host mortality, in both single and coinfection, based on phylogenetic relatedness and the results of the *in vitro* inhibition assays. The survival assay was performed following (Attighi Lorestani *et al*., 2013), with slight modifications. For each single infection or coinfection, five one-day-old age synchronized pupae per replicate were exposed to a conidial suspension of 1x10^8^ spores mL^-1^ in 0.01% Tween 80. Each group of five pupae were immersed in 250µl of spore suspension. After one minute of exposure, the pupae were immediately transferred onto a filter paper and allowed to dry. After 10 minutes and once the pupae started to move, pupae were transferred into a new Petri dish. Coinfections were conducted by preparing spore suspensions at the ratio of 1:1 for each pair of tested fungi. The controls were treated with 0.01% Tween 80 alone. The Petri dishes were covered with a lid and placed in 25 °C and 70±5 humidity. Mortality was recorded for 10 days. Survival was assayed across 11 experimental trials. The design was expanded across trials to cover all combinations of interest, therefore, not all treatments appeared in every trial. Trials 1–4 tested an initial panel of 10 treatments: five single infections (*M. brunneum* KVL 13-06, *M. brunneum* KVL 16-36, *M. lepidiotae, M. pingshaense, M. pemphigi*), an uninfected control, and four co-infections (*M. brunneum* KVL 13-06 + *M. brunneum* KVL 16-36, *M. lepidiotae* + *M. pingshaense, M. lepidiotae* + *M. pemphigi, M. pingshaense* + *M. pemphigi*).

Trials 5–7 tested a set of 8 treatments that introduced the *M. brunneum* KVL 13-06 + *M. pemphigi* co-infection: four single infections (*M. brunneum* KVL 13-06, *M. lepidiotae, M. pingshaense, M. pemphigi*), an uninfected control, and three co-infections (*M. brunneum* KVL 13-06 + *M. pemphigi, M. lepidiotae* + *M. pemphigi, M. pingshaense* + *M. pemphigi*). Trial 8 tested *M. lepidiotae, M. pemphigi*, their co-infection, and an uninfected control. Trial 9 tested *M. brunneum* KVL 13-06, *M. brunneum* KVL 16-36, and *M. pemphigi* as single infections, the *M. brunneum* KVL 13-06 + *M. pemphigi* co-infection, and an uninfected control, and additionally provided the only replicate of the *M. brunneum* KVL 16-36 + *M. pemphigi* co-infection, which was added to complete the set of pairwise co-infections between *M. pemphigi* and each of the other four species. Trial 10 tested *M. pingshaense, M. pemphigi*, their co-infection, and an uninfected control, as previous replicates of this particular paring showed high variation. Trial 11 replicated the combination of *M. lepidiotae, M. pemphigi*, and their co-infection, with an uninfected control. Each individual beetle (*Tribolium castaneum*) was monitored daily for 10 days and represents one data point. This resulted in a final N of 927 individual beetles. Sample sizes are reported per panel of Figure 2 (single 1, single 2, co-infection in each case): A. (*M. lepidiotae* + *M. pingshaense*: 35, 35, 35); B. (*M. lepidiotae* + *M. pemphigi*: 95, 97, 99); C (*M. pingshaense* + *M. pemphigi*: 79, 79, 80); D. (*M. brunneum* KVL 13-06 + *M. pemphigi*: 43, 54, 44); E. (*M. brunneum* KVL 16-36 + *M. pemphigi*: 10, 25, 10); F. (*M. brunneum* KVL 13-06 + *M. brunneum* KVL 16-36: 40, 40, 40).

### Statistical analyses

All analyses were carried out using R-Studio version R version 4.4.1 (2024-06-14 ucrt) (Posit Team, 2023). Radial growth data were analysed using mixed effects linear models comparing the reduction in growth of isolates in combination (growth in combination subtracted from growth alone), using the lmer function from the ‘lme4’ package in R (Bates *et al*., 2015), followed by analysis of deviance tables using the Anova function in the ‘car’ (Fox & Weisberg, 2018) package. The measured isolate and the challenging isolate were included as fixed factors in the models, with trial number and plate number as random factors. Mutliple comparisions were carried out using the emmeans package in R (Searle *et al*., 1980) and p values sidak corrected.

Survival analyses were carried out using a series of pairwise mixed effects Cox proportional hazard models with tested isolate/ combination of isolates as fixed factors and trial number as a random factor, using the ‘coxme’ package in R (Therneau, 2018). Tests were carried out on all available data for each coinfection, along with the corresponding single infections, where these had been tested in parallel. This meant that every dataset was tested at least twice, some more, in order to make all statistical comparisons within each group of all three survival curves. Furthermore, data from single infections were reused in multiple comparisons with differencing coinfection curves, as some trials consisted of multiple strain combinations involving certain isolates. This meant that the single strain comparisons could be compared to multiple coinfection curves. For this reason, all p values were corrected with the fdr method.

## Results

### Radial growth

Overall, all strains showed reduced radial growth in the presence of other *Metarhizium* (Figure 1). This reduction differed significantly depending on the challenging isolate (χ^2^ = 175.03, d.f. = 4, *p* < 0.0001), the measured species (χ^2^ = 126.39, d.f. = 4, *p* < 0.0001), and their interaction (χ^2^ = 313.16, d.f. = 16, *p* < 0.0001) (Figure 1). *M. pemphigi* was by far the strongest inhibitor. As a challenger it significantly reduced the growth of every other isolate (Figure 1; Table 1), consistently producing greater reduction than any other challenger (Figure 1; Table 1). *M. brunneum* 13-06 on the other hand, was significantly less able to inhibit other *Metarhizium* than the other isolates tested, particularly when challenging *M. lepidiotae* and *M. pingshaense* (Table 1; Figure 1). The tested isolates also differed in how inhibited they were by other *Metarhizium. M. pingshaense* was the least sensitive to inhibition, showing lower reduction in growth than any other isolate (Figure 1).

**Table 1.**
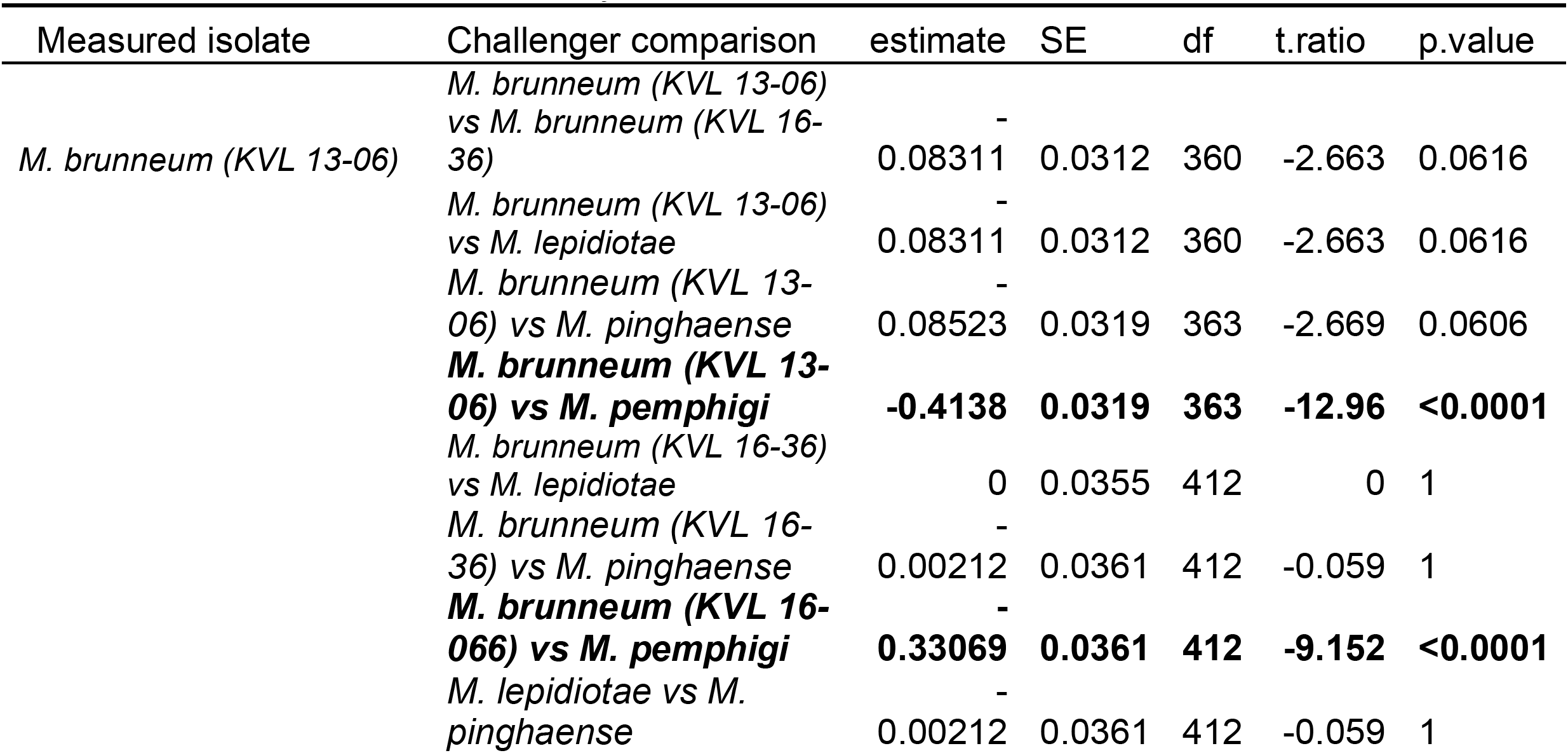
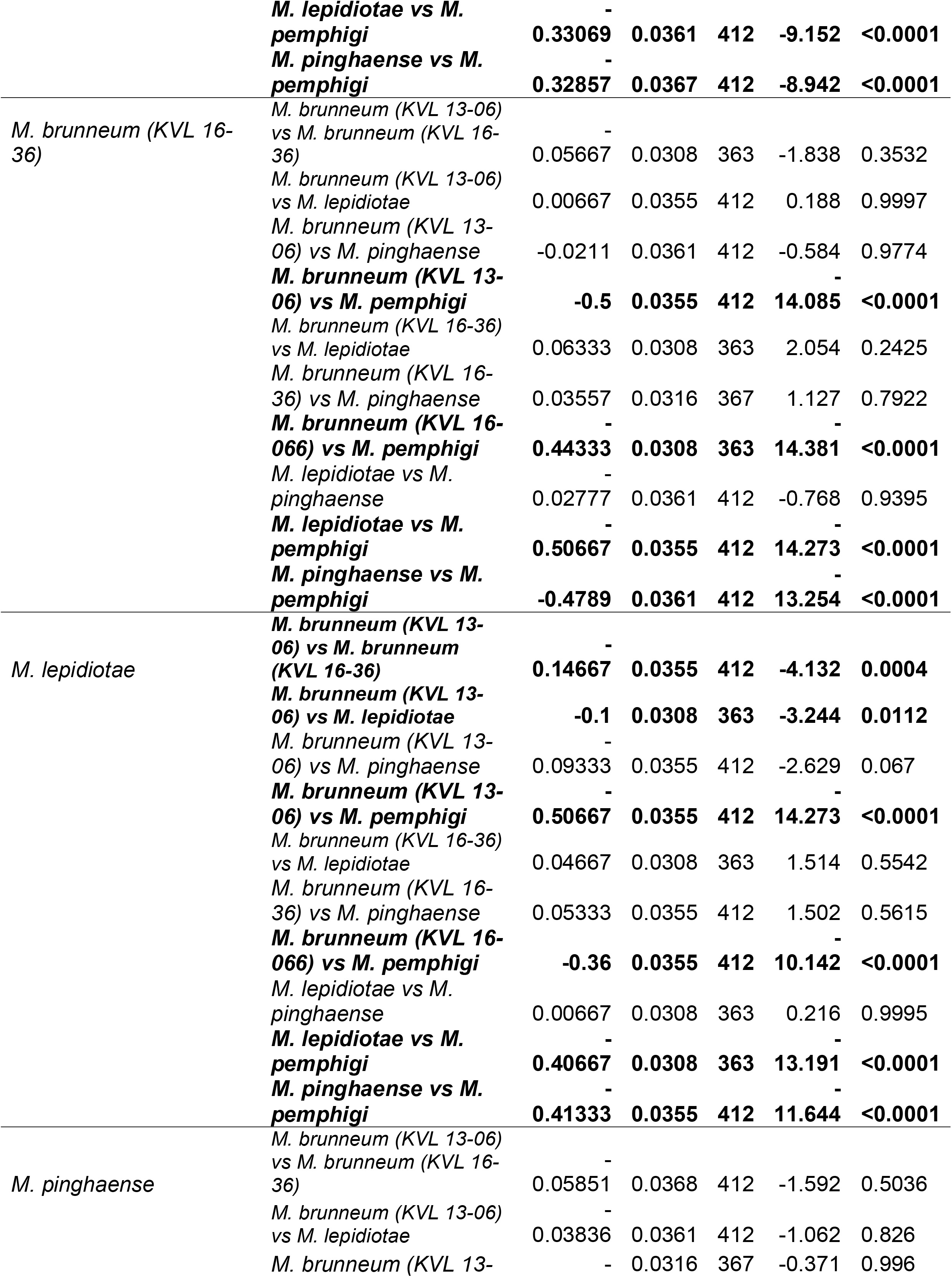
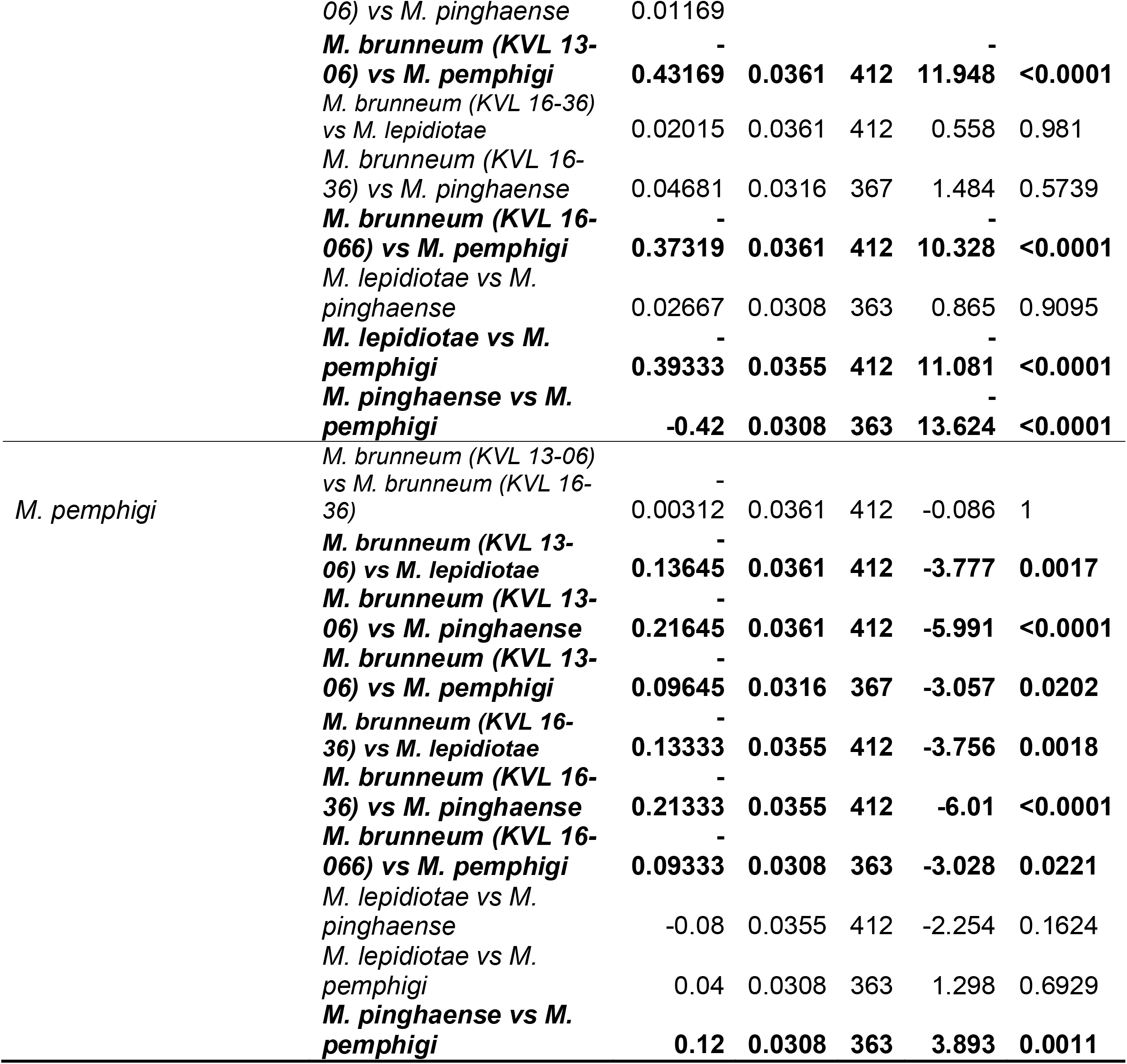
Multiple comparisons of growth reduction among challenged isolates within each measured isolate. P values are sidak adjusted.

**Figure 1.**
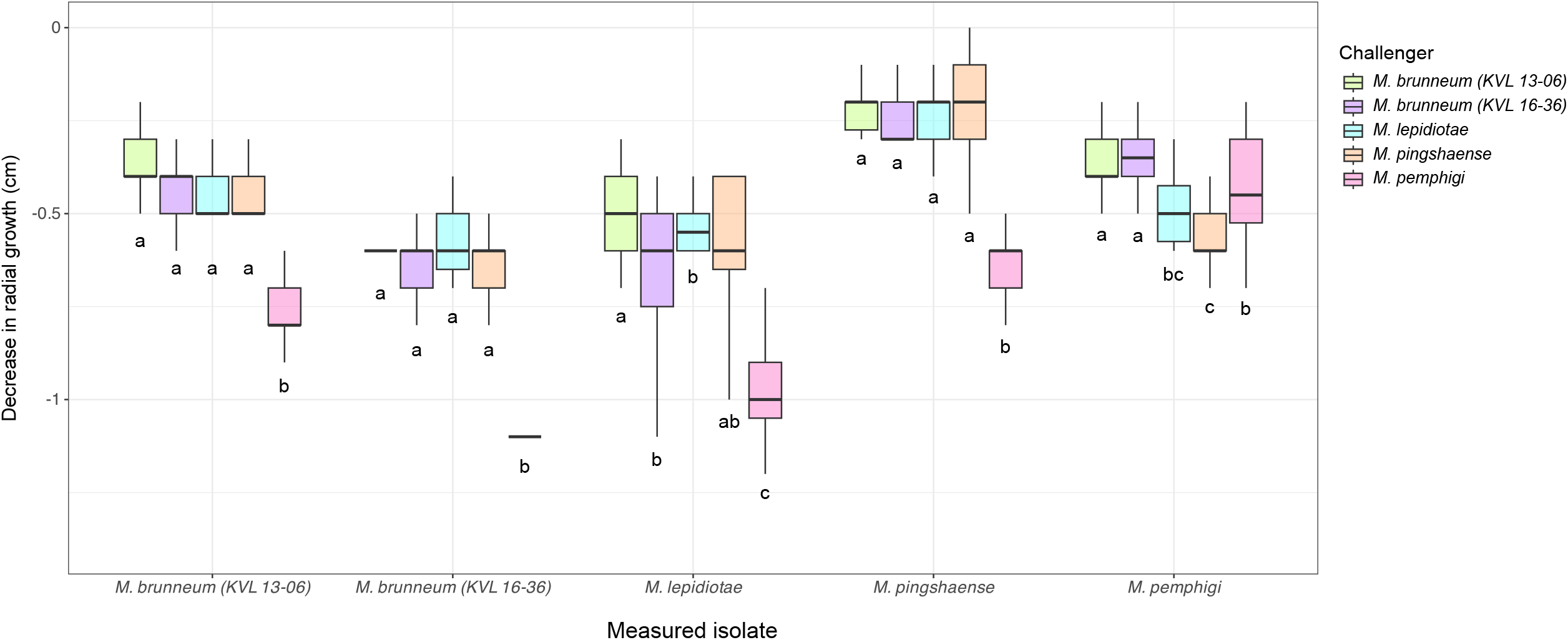
Difference in radial growth between each pair of isolates calculated as the difference in diameter when grown in co-culture in comparison to diameter when grown alone.

### Survival analyses

To test how coinfection impacted host survival, we carried out survival analysis comparing pairs of isolates in single and coinfections. We tested the single and combined effects of: *M. lepidiotae* and *M. pingshaense, M. lepidiotae* and *M. pemphigi, M. pingshaense* and *M. pemphigi, M. brunneum* KVL 13-06 and *M. pemphigi, M. brunneum* KVL 16-36 and *M. pemphigi* and *M. brunneum* KVL 13-06 and *M. brunneum* KVL 16-36. *M. pemphigi* was tested against all other species due to its strong inhibitory effects and the fact that it is the most phylogenetically distant from all other species, belonging to the flavovride clade of *Metarhizium* (Glare *et al*., 2021; Kobmoo *et al*., 2024). The remaining combinations represented different levels of phylogenetical relatedness, *M. lepidotiae* and *M. pighaense* belong to the sister clades MGB and PARB respectively (Glare et al. 2021; Kobmoo et al. 2024) and the two *Metarhizium brunneum* strains belong to the same species. We only observed significant differences in the combinations involving *M. lepidiotae* and *M. pingshaense* and *M. brunneum* KVL 13-06 and *M. pemphigi* (Figure 2; Table 2). Single infections did not differ significantly from one another or from coinfection in any of the other combinations (Figure 2; Table 2). In the case of *M. lepidiotae* and *M. pingshaense*, virulence differed significantly between single infections and coinfections. *M. lepidiotae* caused faster host death than *M. pingshaense* (coef = 1.021, Hazard Ratio = 2.777, z = 2.79, *p* = 0.03144). In co-infection, killing speed did not differ significantly from that of *M. pingshaense in* single infection (coef = −0.018, HR = 0.982, z = −0.04, *p* = 0.965) but killing was significantly slower than by *M. lepidiotae* in single infection (coef = −1.021, HR = 0.360, z = −2.79, *p* = 0.03144) (Figure 2A). In the case of *M. brunneum* KVL 13-06 and *M. pemphigi*, co-infection resulted in significantly lower host mortality than in single infection with *M. pemphigi* (coef = −0.798, HR = 0.450, z = −2.91, *p* = 0.03144) (Figure 2D), with coinfection resulting in lower virulence than either single infection.

**Table 2.** Results of pairwise survival analyses with raw and fdr corrected p values. All comparisons were carried out using mixed effects cox proportional hazard models with isolate combination as a fixed factor and trial number as a random factor.

| Group 1 | Group 2 | raw p value | p value (fdr) |
| --- | --- | --- | --- |
| *Metarhizium pemphigi* | ***brunneum 13-06+pemphigi*** | **0.00357** | **0.03144** |
| *Metarhizium lepidiotae* | ***lepidiotae+pinghaense*** | **0.0045** | **0.03144** |
| *Metarhizium lepidiotae* | ***Metarhizium pingshaense*** | **0.00524** | **0.03144** |
| *Metarhizium brunneum* KVL 16-36 | *Metarhizium pemphigi* | 0.0183 | 0.08235 |
| *Metarhizium brunneum* KVL 16-36 | *Metarhizium brunneum* KVL 13-06 | 0.0283 | 0.10188 |
| *Metarhizium pingshaense* | *Metarhizium pemphigi* | 0.0376 | 0.1128 |
| *Metarhizium brunneum* KVL<br>13-06 | *brunneum 13-06+pemphigi* | 0.048 | 0.123428571 |
| *Metarhizium pemphigi* | *pinghaense+pemphigi* | 0.1063 | 0.239175 |
| *Metarhizium brunneum* KVL<br>13-06 | *brunneum 13-06+16-36* | 0.1712 | 0.3424 |
| *Metarhizium brunneum* KVL<br>16-36 | *brunneum 16-36+pemphigi* | 0.2294 | 0.41292 |
| *Metarhizium pemphigi* | *lepidiotae+pemphigi* | 0.263 | 0.430363636 |
| *Metarhizium brunneum* KVL<br>13-06 | *Metarhizium pemphigi* | 0.37752 | 0.56628 |
| *Metarhizium lepidiotae* | *Metarhizium pemphigi* | 0.478 | 0.5736 |
| *Metarhizium brunneum* KVL<br>16-36 | *brunneum 13-06+16-36* | 0.435 | 0.5736 |
| *Metarhizium pemphigi* | *brunneum 16-36+pemphigi* | 0.4532 | 0.5736 |
| *Metarhizium pingshaense* | *pinghaense+pemphigi* | 0.6187 | 0.6960375 |
| *Metarhizium lepidiotae* | *lepidiotae+pemphigi* | 0.706 | 0.747529412 |
| *Metarhizium pingshaense* | *lepidiotae+pinghaense* | 0.96506 | 0.96506 |

**Figure 2.**
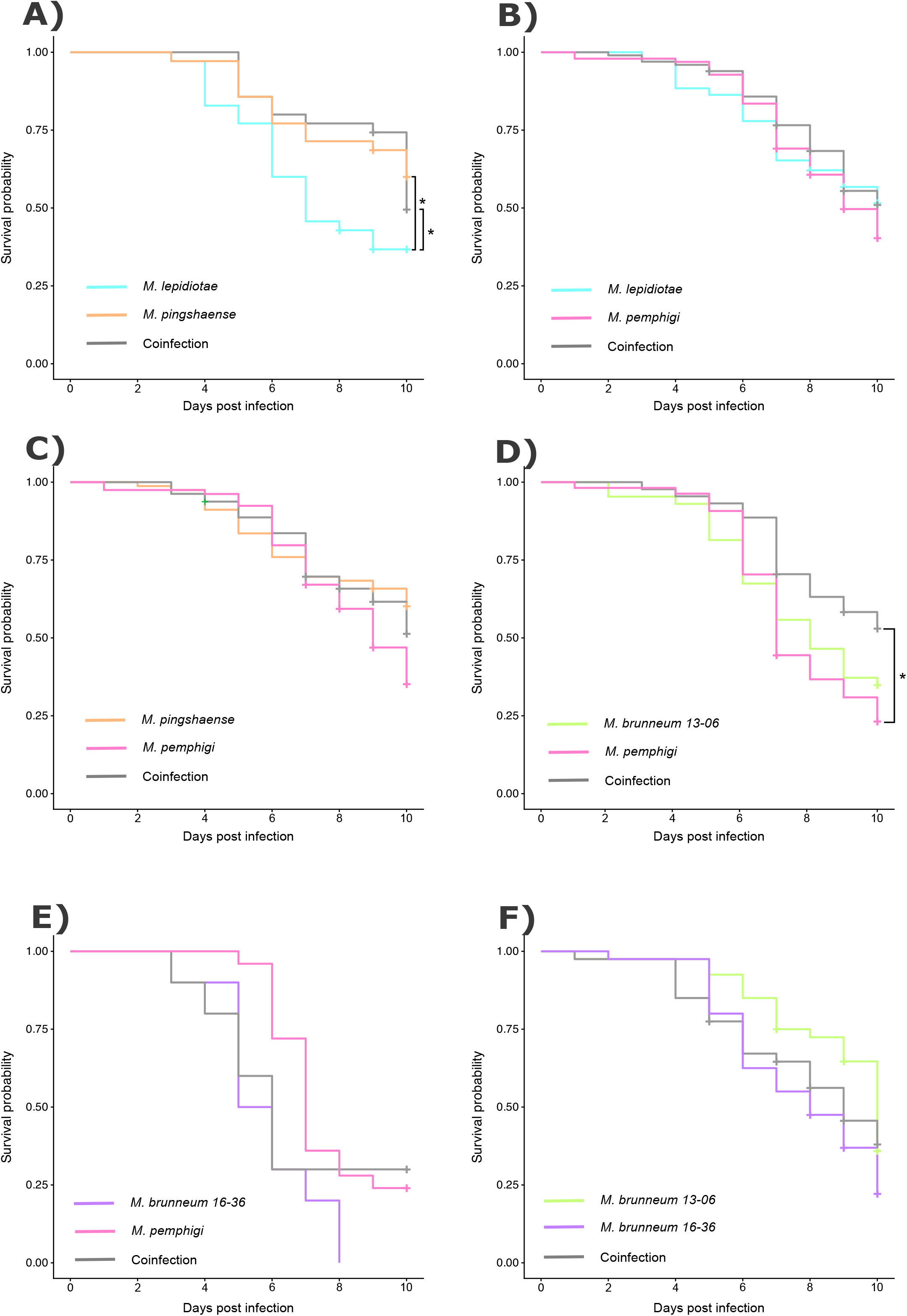
Kaplan–Meier survival curves of *T. castaneum* pupae following *Metarhizium* topical immersion of selected pairs of isolates in single and co-infections. Survival curves were analyzed using Kaplan–Meier survival analysis followed by cox proportional hazard test A) *M*.*lepidiotae* and *M. pingshaense* infecting alone and in coinfection; B) *M. pemphigi* and *M. lepidiotae* infecting alone and coinfection; C) *M. pingshaense* and *M. pemphigi* infecting alone and coinfection; D) *M. brunneum* KVL 13-06 and *M. pemphigi* infecting alone and coinfection; E) *M. brunneum* KVL 16-36 and *M. pemphigi* infecting alone and coinfection; F) *M. brunneum* KVL 13-06 and *M. brunneum* KVL 16-36 infecting alone and coinfection.

## Discussion

We found that isolates were inhibited by and inhibitory towards one another to differing degrees, and that this inhibition was to an extent but not completely reflected in the virulence outcomes of the strain pairs in coinfection. *M. pingshaense* was the strain least susceptible to inhibition by other fungi and indeed, in the only coinfection where single infection differed significantly from a coinfecting partner, *M. lepidiota*e, with the survival curve of infected *T. castaneum* reflecting that of *M. pingshaense*. Here the overall virulence of the coinfection was reduced in comparison to single infection with the more virulent parasite. The most inhibitory isolate of all, *M. pemphigi*, did not significantly differ in virulence compared to any strain that it was paired with in coinfection, however, in coinfection with *M. brunneum* 13-06, virulence was lower than that of either of the coinfecting partners in single infection.

Interestingly, *M. pemphigi* was able to inhibit other *Metarhizium* strains to a much great extent than any other tested strain. The fact that *M. pemphigi* is able to inhibit all other strains while avoiding the same degree of self-inhibition, may be a consequence of the phylogenetic relationship among tested strains. *M. pemphigi* is phylogenetically distinct from all other tested strains, belonging to the flavoride clade of, *Metarhizium*, which was the first to diverge from the rest of the group (Glare *et al*., 2021; Kobmoo *et al*., 2024). This phylogenetic distinction may mean that *M. pemphigi* is different enough from the strains, that it is able to produce inhibitory secondary metabolites that it itself is not inhibited by. To our knowledge, little is known about the anti-fungal antibiotics and secondary metabolites secreted my *M. pemphigi*, however, the production of secondary metabolites in fungi is common, often facilitating interference competition (Chatterjee *et al*., 2016; Künzler, 2018). At least in the absence of a host, these secretions seem to be critical in mediating interactions with other fungi.

Although relatedness, may have impacted our *in vitro* inhibition results, it did not have a clear impact on virulence. In general, theory predicts that kin selection should lead to cooperation in coinfections of closely related strains, whereas infections with more distantly related parasites may lead to rapid growth as a consequence of competition and thus higher virulence (Alizon *et al*., 2013; Buckling *et al*., 2007). Specifically, whether coinfecting strains are strains of the same or different species is predicted to influence virulence in coinfection (Alizon *et al*. 2013). Species concepts are complicated in fungi, and species assignments in *Metarhizium* have been challenging (Kobmoo *et al*., 2024; Schuster *et al*., 2023, 2026), nevertheless according to the current phylogeny, our experimental tests included pairs where we tested two strains of the same species, two species of the same clade, two species of sister clades, along with pairs of more distantly related species. Despite the high inhibitory capacity of *M. pemphigi* towards the other fungal species, however, its presence does not universally impact virulence.

In general, we saw similar virulence levels among strains, which could explain why there were only significant differences in two tested combinations in coinfection. Interestingly, however, in coinfection between *M. pemphigi* and *M. brunneum*, overall virulence of the coinfection was significantly lowered compared to single infection with *M. pemphigi*, this was in spite of both strains showing similar virulence levels in single infection. The type of competition dynamics between colonising microbes is known to impact virulence outcomes, with interference competition leading to virulence reductions in coinfection (Dutt *et al*., 2022; Vorburger & Perlman, 2018). For example, virulence in coinfections of *Galleria mellonella* larvae with bacteriocin producing bacteria was lower than in either single infection in cases where bacteria where able to inhibit one another *in vitro* (Massey *et al*., 2004). A similar pattern is also seen over evolutionary time in *Bacillus thuringiensis*, where strains evolving in multi-strain coinfections evolve increased competitiveness at the cost of virulence (Garbutt *et al*., 2011). Indeed, this pattern is mirrored in our data from coinfection of *M. lepidiotae* and *M. pingshaense*. Overall *M. pigsaense* was the strain least inhibited by others. In the one case where *M. pingshaense* differed in virulence from its coinfecting partner, in this case *M. lepidiotae*, virulence mirrored that of *M. pingsanse alone*, the less virulent and least susceptible to inhibition of the two. Our findings thus provide some tentative empirical support for theoretical predictions that parasites in coinfections face allocation trade-offs between investing in competition with other parasites and exploiting the host for growth and transmission (Alizon *et al*., 2013; Mideo, 2009; Van Baalen & Sabelis, 1995)

Overall, in the cases where we saw significant differences between coinfections and individual single infections, coinfections were less virulent than single infections. The most common outcome of coinfection is in fact that virulence mirrors that of the more virulent parasite, but with substantial variation and certainly not without exceptions (Rafaluk *et al*., 2026). This is a pattern seen across many bacterial coinfections (Schmitz *et al*., 2023). The reason that our results here differ from this general trend is likely a consequence of interference competition. When parasites directly attack one another, virulence tends to be reduced (Garbutt *et al*., 2011; Massey *et al*., 2004). In addition, our results are consistent with a previous study of *Metarhizium* coinfections. Sheng and St Ledger (2024) found that in coinfections of both *Dropsophila melanogaster* and *Manducta sexta* with *Metarhizium robertsii* and *Metarhizium anisopliae*, virulence is lower in coinfection than single infections and that there was spatial separation in these coinfections (Sheng & St. Leger, 2024). These results together with ours suggest that interference competition is likely to be key in driving virulence evolution in *Metarhizium*.

## Conclusions

Understanding the interaction dynamics of coinfecting parasites is crucial for disease management, as well as understanding the ecology and evolution of infections. Here we show in a species with high levels of strain – strain inhibition, that interference competition at least to an extent influenced virulence outcomes in coinfection, reducing the overall virulence of the infection. This is consistent with findings from other biological systems where interference competition or putative interference competition has reduced virulence in coinfection (Garbutt *et al*., 2011; Massey *et al*., 2004; Sheng & St. Leger, 2024). These results together with ours highlight the fact that, while in general virulence in coinfection may be determined by the more virulence parasite (Rafaluk *et al*., 2026; Schmitz *et al*., 2023), in systems with interference competition, coinfection can result, in fact, in reduced virulence.

## Ethics

n/a

## Author contributions

N.G.: investigation, formal analysis, writing original draft; C.O.: investigation, formal analysis;

N. K. investigation, S. K. investigation, C.R.: supervision, conceptualisation, formal analysis, writing, review and editing. All authors gave final approval for publication.

## Declaration on use of AI

We did not use any ai in the preparation of this manuscript or analysis.

## Acknowledgements

Thanks to Henrik Hjarvard de Fine Licht, Nicolai Meyling, and Dino McMahon for providing the fungal isolates used in this experiment and to Joachim Kurtz for providing the beetles. Thanks to Elisa Bittermann, Fabian Günther, and Carlotta Eisenhut for help and support in the lab Mathias Franz for statistical advice, and Caroline Zanchi and Maryam Keshavarz for helpful advice on the lab work and infection protocol. Thanks to Jens Rolff for general support and feedback on an earlier version of the manuscript.

