## Supplementary material for "Interference competition influences virulence in *Metarhizium* coinfections": S1

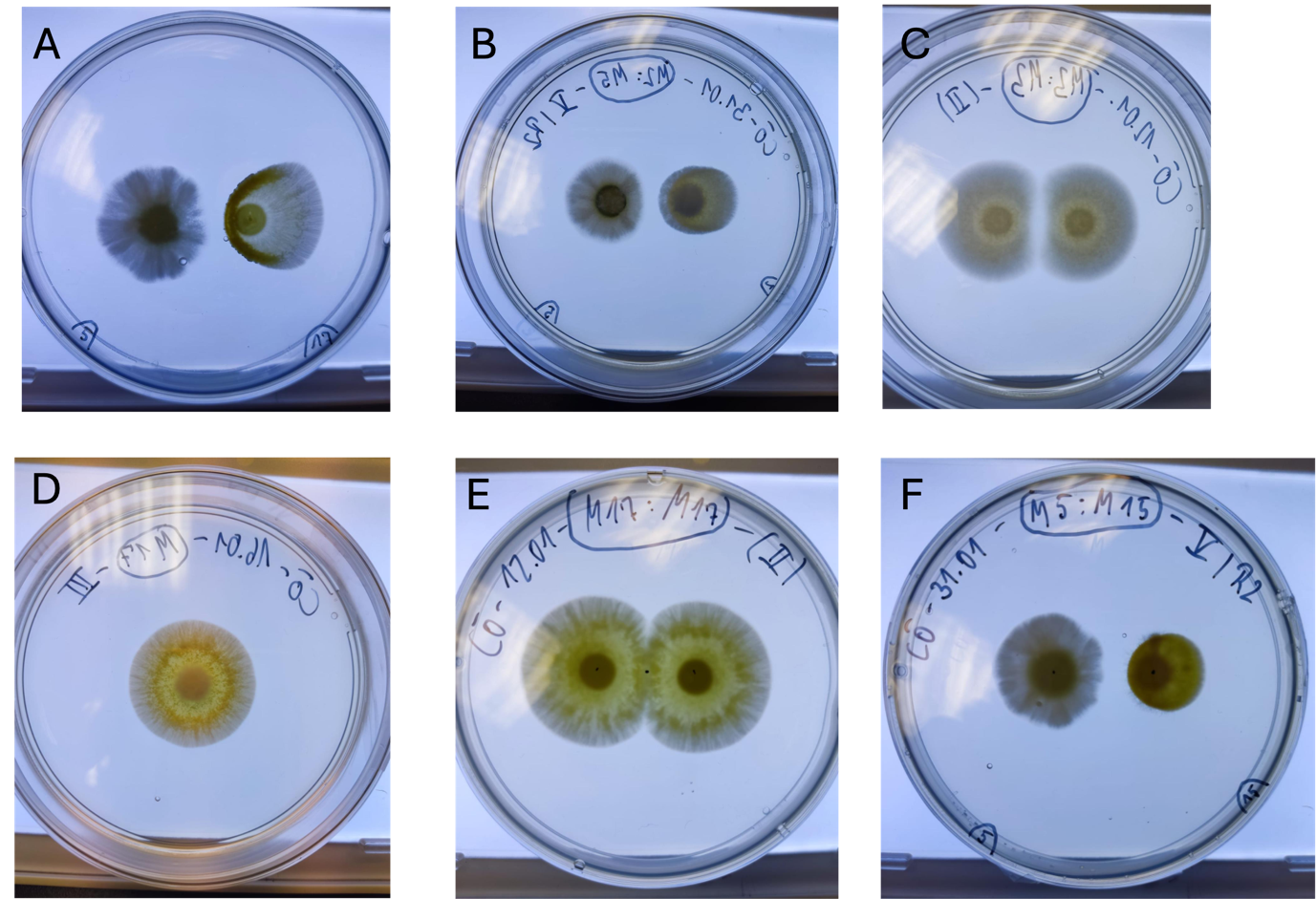

Figure S1. Representative plates of different strain combinations, showing the roughly spherical colony form. A) *M. pemphigi* and *M. brunneum* KVL 16-36; B) *M. lepidiotae* and *M. pemphigi*; C) *M. pingshaense*; D) and E) M. *brunneum* strain KVL 16-36 F) *M. pemphigi* and *M. brunneum* strain KVL 13-06.
